# powerEQTL: An R package and shiny application for sample size and power calculation of bulk tissue and single-cell eQTL analysis

**DOI:** 10.1101/2020.12.15.422954

**Authors:** Xianjun Dong, Xiaoqi Li, Tzuu-Wang Chang, Scott T. Weiss, Weiliang Qiu

**Affiliations:** Genomics and Bioinformatics Hub, Brigham and Women’s Hospital, Harvard Medical School; Precision Neurology Program, Brigham and Women’s Hospital, Harvard Medical School; Molecular Pathological Epidemiology Program, Brigham and Women’s Hospital, Harvard Medical School; Channing Division of Network Medicine, Brigham and Women’s Hospital, Harvard Medical School; Non-Clinical Efficacy & Safety, Biostatistics & Programming, Sanofi

**Author notes:** These authors contributed equally (Xianjun Dong, Xiaoqi Li).

## Abstract

**Summary:** Genome-wide association studies (GWAS) have revealed thousands of genetic loci for common diseases. One of the main challenges in the post-GWAS era is to understand the causality of the genetic variants. Expression quantitative trait locus (eQTL) analysis has been proven to be an effective way to address this question by examining the relationship between gene expression and genetic variation in a sufficiently powered cohort. However, it is often tricky to determine the sample size at which a variant with a specific allele frequency will be detected to associate with gene expression with sufficient power. This is particularly demanding with single-cell RNAseq studies. Therefore, a user-friendly tool to perform power analysis for eQTL at both bulk tissue and single-cell level will be critical. Here, we presented an R package called powerEQTL with flexible functions to calculate power, minimal sample size, or detectable minor allele frequency in both bulk tissue and single-cell eQTL analysis. A user-friendly, program-free web application is also provided, allowing customers to calculate and visualize the parameters interactively.

**Availability and implementation:** The powerEQTL R package source code and online tutorial are freely available at CRAN: https://cran.r-project.org/web/packages/powerEQTL/. The R shiny application is publicly hosted at https://bwhbioinfo.shinyapps.io/powerEQTL/.

**Contact:** Xianjun Dong, Weiliang Qiu

**Supplementary information:** Supplementary data are available at Bioinformatics online.

## Introduction

Genome-wide association studies (GWAS) have revealed genetic risk loci for thousands of traits or diseases (MacArthur *et al*., 2017; Buniello *et al*., 2019). Nearly 90% of the GWAS loci are located in non-coding regions (Edwards *et al*., 2013), suggesting that they may play a role by influencing gene expression. One of the main challenges in the post-GWAS era is understanding how these genetic variants cause the phenotype, for example, by regulating the expression of disease-associated or tissue-specific genes. Expression quantitative trait locus (eQTL) analysis has provided such a framework to test the effect of genetic variation on gene expression (Nica and Dermitzakis, 2013). For instance, the Genotype-Tissue Expression (GTEx) project has performed eQTL analysis between genetic variation and genome-wide gene expression in 54 non-diseased tissue sites across nearly 1000 individuals, providing a comprehensive public resource to understand the effect of genetic variants in a wide spectrum of tissue bank samples (GTEx Consortium, 2013, 2015). Enhancing GTEx (eGTEx) further extended this effort to include more intermediate molecular phenotypes other than gene expression (eGTEx Project, 2017). Recent increases in single-cell genomics will allow mapping eQTLs across different cell types, in dynamic processes, and in 3D spaces, many of which are obscured when using bulk methods (van der Wijst *et al*., 2018, 2020). One of the critical steps common to all eQTL experiments is to determine the minimum sample size with enough power to detect variants with a low frequency (e.g., minor allele frequency less than 5%) but a substantial effect on gene expression. However, there is no such tool available for sample size and power calculation for eQTL analysis.

Here we developed equation-based statistical models to calculate sample size and power for an eQTL analysis in both bulk tissue and single-cell settings. The tool, called powerEQTL, was implemented in both an R package and an interactive online application.

## Materials and methods

### Bulk tissue eQTL

Bulk tissue eQTL is to identify the downstream effects of disease-associated genetic variants on the gene expression measured at the bulk tissue level. Because of the affordable price (compared to a single-cell experiment) and the convenience to get enough volume of RNAs from bulk tissue, bulk RNA-sequencing is still the most widely used technique to profile the transcriptome of a tissue nowadays. Gene expression values were quantified on tissue homogenates, usually one sample per subject, for a number of subjects. Normalized gene expressions were then compared among groups of subjects with different genotypes. Since the effect sizes of eQTL are usually small and the large number of gene-SNP pairs leads to a multiple-testing issue (Huang *et al*., 2018), a proper power analysis including sample size and power calculation is needed before performing experiments.

We implemented the power analysis of bulk tissue eQTL based on two different models, one-way unbalanced ANOVA and simple linear regression (see Online Supplementary Document). They both test for the potential association between genotype and gene expression. The difference lies in that ANOVA test treats the genotype as a categorical data (e.g., AA, AB, and BB) and tests the potential non-linear association, while simple linear regression treats genotype as continuous variable using additive coding (e.g., 0 for AA, 1 for AB, and 2 for BB, where B is the minor allele) and tests the linear association. GTEx project used the one-way unbalanced ANOVA model in their analysis (GTEx Consortium, 2013). We implemented the two models in functions of *powerEQTL*.*ANOVA* and *powerEQTL*.*SLR* in our R package, respectively. Note that if we know the association is linear, *powerEQTL*.*SLR* would be more powerful than *powerEQTL*.*ANOVA*. This is because categorizing a continuous-type variable to a set of nominal-type variables would lose information.

Since type I error rate (α), type II error rate (β or 1-power), effect size, and sample size are interrelated in power analysis, we could calculate any one of them if we know the rest three. So, we also implemented the functions like that, e.g., calculating one of the 4 parameters (power, sample size, slope, and minimum allowable MAF) by setting the corresponding parameter as NULL and providing values for the other three parameters in *powerEQTL*.*SLR*.

### Single-cell eQTL

Unlike bulk tissue RNAseq, single-cell RNAseq usually profiles thousands of cells per sample, which provides a better representation for the gene expression distribution of a tissue than a single value from bulk RNAseq. However, the gene expressions among cells within a sample are not independent, e.g., cells from one tissue sample are assumed more correlated than cells between samples. The structured data requires a different model for power analysis. We implemented the single-cell eQTL by modeling the single-cell RNA expression and genotype in a linear mixed effects model: y_ij_ =β_0i_ + β_1_*x_i_ + ε_ij_, where y_ij_ is the gene expression level for the j^th^ cell of the i^th^ subject, x_i_ is the genotype for the i^th^ subject using additive coding (e.g., 0, 1, and 2. See Online Supplementary Document for details). The power to test if the slope β_1_ is equal to zero is implemented in the function *powerEQTL*.*scRNAseq* with parameters of subject size (*n*), number of cells per subject (*m*), slope (β_*1*_), standard deviation of the gene expression (σ_*y*_), MAF, intra-subject correlation (i.e., correlation between y_ij_ and y_ik_ for the j^th^ and k^th^ cells of the i^th^ subject, ρ), and number of SNP-gene pairs (*nTest*). Similarly, the function can be used to calculate one of the 4 parameters (power, sample size, minimum detectable slope, and minimum allowable MAF) by setting the corresponding parameter as NULL and providing values for the other 3 parameters.

## Result

The powerEQTL R package is available in CRAN and has been downloaded over 10,000 times since its first deployment (see Figure 1). We also implemented the functions for power and sample size calculation in an online, interactive, program-free web application using R Shiny. Power curves of different MAFs for multiple sample sizes are visualized and downloadable for both bulk tissue and single-cell eQTL. The calculator pages allow users to freely play with the parameters for tissue and single-cell eQTL power analysis.

**Figure 1.**
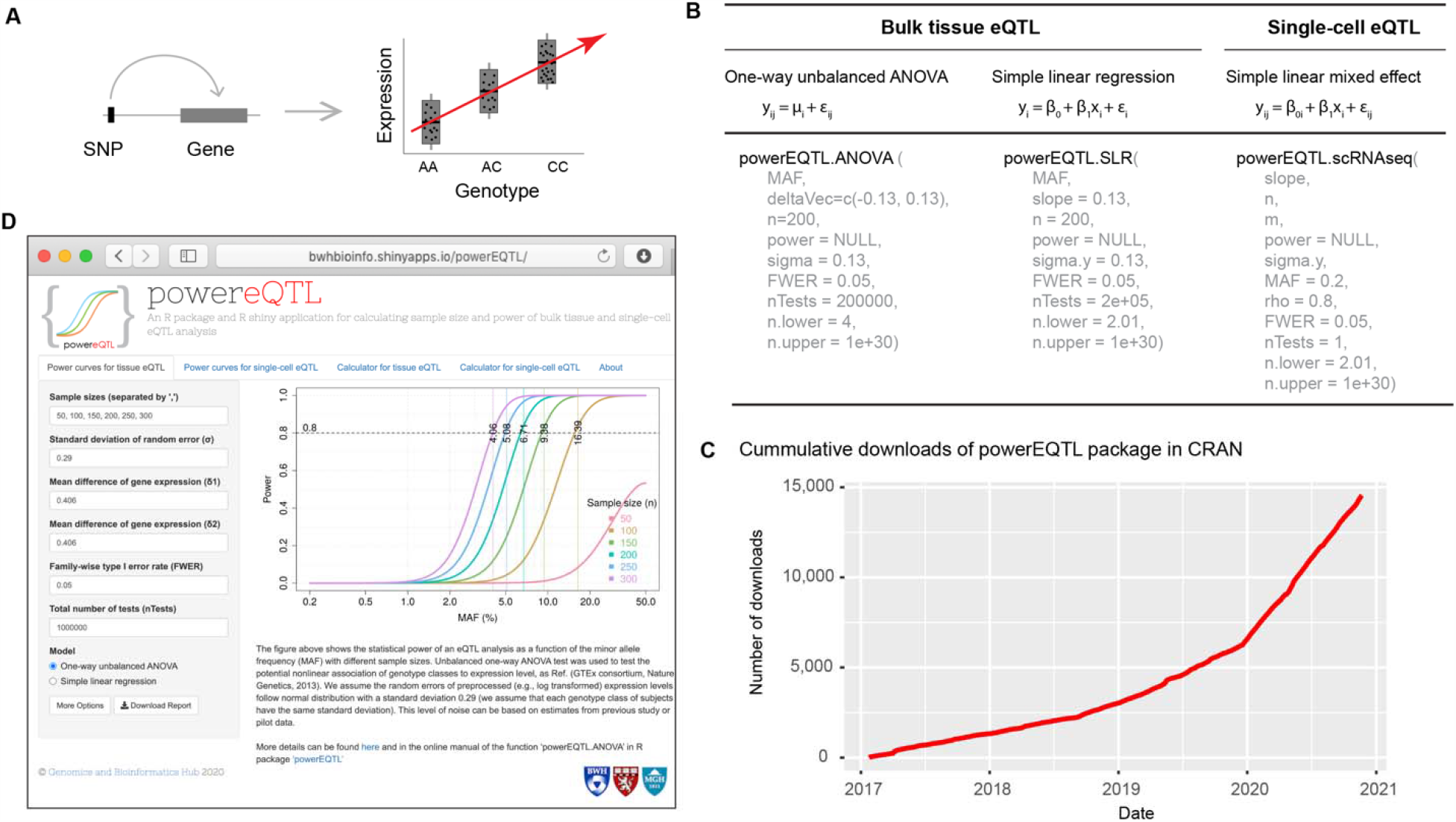
(A) eQTL schema. (B) The main models and functions in the powerEQTL package. (C) Downloads summary of powerEQTL since its original repository on CRAN (data generated by cranlog R package). (D) Screenshot of powerEQTL R shiny application.

## Discussion

While several R or Bioconductor packages are available for omics sample size and power calculation, such as sizepower (equation-based, 2006), RNASeqPower (equation-based, 2013), PROPER (simulation-based, 2015), powsimR (simulation-based, 2017), RnaSeqSampleSize (2018), ssizeRNA (equation-based, 2019), PowerSampleSize, pwrEWAS, and powerGWASinteraction, none are for eQTL analysis yet that we are aware of. To apply powerEQTL to RNAseq data, appropriate data transformation, such as voom (Law *et al*., 2014) and countTransformers (Zhang *et al*., 2019), is needed to convert counts to continuous data. In addition to scRNAseq, other structured data, such as scATACseq, single-cell methylation, grouped cell lines etc. can also be applied to this eQTL model.

