## Supplementary material for "powerEQTL: An R package and shiny application for sample size and power calculation of bulk tissue and single-cell eQTL analysis": Online Supplementary Document

**A Vector representation 15**

**B Generalized least squares estimate when variance-covariance matrix is known 17**

**C Mean, variance, and distribution of the generalized least squares estimate 18**

**D Calculating the power for testing if the slope is equal to zero 18**

**E Calculation of the variance of the slope estimate 20**

**F Power calculation formula revisit 24**

**G Variance of genotype under Hardy-Weinberg Equilibrium 24**

**H Power calculation formula for genotypes under Hardy-Weinberg Equilibrium 25**

### 1 Power Calculation for Bulk Tissue eQTL Based on ANOVA

#### 1.1 Introduction

If we would like to test potential non-linear relationship between genotype of a SNP and expression of a gene, we can use un-balanced one-way ANOVA. Actually, an article published by the GTEx Consortium in 2013[3] used this approach.

#### 1.2 General formula

Suppose there are $k = 3$ groups of subjects: (1) mutation homozygotes; (2) heterozygotes; and (3) wildtype homozygotes. We would like to test if the mean expression $\mu_i$, $i = 1, \ldots, k$, of the gene is the same among the $k$ groups of subjects. We can use the following one-way ANOVA model to characterize the relationship between observed gene expression level $y_{ij}$ and the population mean expression level $\mu_i$:

$$\begin{aligned} y_{ij} =& \mu_i + \epsilon_{ij}, \quad \epsilon_{ij} \sim N\left(0, \sigma^2\right), \\ i =& 1, \ldots, k, \\ j =& 1, \ldots, n_i, \end{aligned} \tag{1}$$

where $y_{ij}$ is the observed gene expression level for the $j$-th subject in the $i$-th group, $\mu_i$ is the mean gene expression level of the $i$-th group, $\epsilon_{ij}$ is the random error, $k$ is the number of groups, $n_i$ is the number of subjects in the $i$-th group. Denote the total number of subjects as $N = \sum_{i=1}^{k} n_i$. That is, we have $n_1$ mutation homozygotes, $n_2$ heterozygotes, and $n_3$ wildtype homozygotes.

We would like to test the null hypothesis $H_0$ and alternative hypothesis $H_1$:

$$\begin{aligned} &H_0 : \mu_1 = \mu_2 = \mu_3, \\ &H_1 : \text{not all } \mu_i \text{ are the same.} \end{aligned} \tag{2}$$

It is well know that the test statistic for Hypotheses (2) is the F statistic

$$F = \frac{MS_{grp}}{MSE}, \tag{3}$$

where $MS_{grp}$ is the mean square of group effect and $MSE$ is the mean square of error.

We will reject the null hypothesis $H_0$ if the test statistic $F$ is large enough. The type I error rate $\alpha$ is defined as

$$\alpha = Pr\left(F > c | H_0\right) = 1 - Pr\left(F < c | H_0\right), \tag{4}$$

where $c$ is the rejection boundary (i.e., cutoff).

Under $H_0$, the test statistic $F$ follows the F distribution with degrees of freedom $df_1 = k - 1$ and $df_2 = N - k$ (Denote it as $F_{k-1,N-k}$). Hence, the cutoff $c$ is the upper $100\alpha$

percentile of the F distribution $F_{k-1,N-k}$. Denote

$$c = F_{1-\alpha}\left(k-1, N-k\right). \tag{5}$$

Under $H_1$, the test statistic $F$ follows the non-central F distribution with degrees of freedom $df_1 = k-1$, $df_2 = N-k$, and non-centrality parameter $\lambda$ (Denote it as $F_{k-1,N-k,\lambda}$).

According to O'Brien and Muller (1993)[2], the non-central parameter $\lambda$ is defined as

$$\lambda = \frac{N}{\sigma^2}\sum_{i=1}^{k}\omega_i\left(\mu_i-\mu\right)^2, \tag{6}$$

where $\mu_i$ is the mean value for the $i$-th level, $\omega_i = n_i/N$ is the weight for the $i$-th level, and $\mu$ is the overall mean,

$$\mu = \sum_{i=1}^{k}\omega_i\mu_i. \tag{7}$$

Hence, the power calculation formula for un-balanced one-way ANOVA with $k$ levels is

$$\begin{aligned} power =& Pr\left(F > c|H_1\right) \\ =& Pr\left(F \geq F_{1-\alpha}\left(k-1, N-k\right)\middle| F \sim F_{k-1,N-k,\lambda}\right)\right). \end{aligned} \tag{8}$$

#### 1.3 Simplification of non-central parameter

For our case, $k = 3$ and the non-centrality parameter $\lambda$ can be rewritten as

$$\lambda = \frac{N}{\sigma^2}\sum_{i=1}^{3}w_i\left(\mu_i-\mu\right)^2. \tag{9}$$

By assuming Hardy-Weinberg Equilibrium, we have

$$\begin{aligned} w_1 &= \frac{n_1}{N} = \theta^2 \text{ (genotype frequency for mutation homozygotes)}, \\ w_2 &= \frac{n_2}{N} = 2\theta(1-\theta) \text{ (genotype frequency for heterozygotes)}, \\ w_3 &= \frac{n_3}{N} = (1-\theta)^2 \text{ (genotype frequency for wildtype homozygotes)}, \end{aligned}$$

where $\theta$ is the minor allele frequency (MAF).

We assume the mean gene expression levels for the 3 groups are

$$
\begin{aligned}
&\mu_1 = a - \delta_1 \text{ (for mutation homozygotes)},\\
&\mu_2 = a \text{ (for heterozygotes)},\\
&\mu_3 = a + \delta_2 \text{ (for wildtype homozygote)}.
\end{aligned}
$$

That is,

$$
\begin{aligned}
\mu_2 - \mu_1 &= \delta_1,\\
\mu_3 - \mu_2 &= \delta_2.
\end{aligned}
$$

Next, we would like to show that the non-centrality parameter $\lambda$ depends only on $\delta_1$ and $\delta_2$, but not depends on $a$.

We can obtain

$$
\begin{aligned}
\mu =&\omega_1\mu_1 + \omega_2\mu_2 + \omega_3\mu_3\\
=&\frac{n_1}{N}(a-\delta_1) + \frac{n_2}{N}a + \frac{n_3}{N}(a+\delta_2)\\
=&a\left(\frac{n_1}{N} + \frac{n_2}{N} + \frac{n_3}{N}\right) - \frac{n_1}{N}\delta_1 + \frac{n_3}{N}\delta_2\\
=&a - \frac{n_1}{N}\delta_1 + \frac{n_3}{N}\delta_2.
\end{aligned}
$$

Hence,

$$
\begin{aligned}
\mu_1 - \mu =&(a-\delta_1) - \left(a - \frac{n_1}{N}\delta_1 + \frac{n_3}{N}\delta_2\right)\\
=&-\delta_1 + \frac{n_1}{N}\delta_1 - \frac{n_3}{N}\delta_2\\
\mu_2 - \mu =&a - \left(a - \frac{n_1}{N}\delta_1 + \frac{n_3}{N}\delta_2\right)\\
=&\frac{n_1}{N}\delta_1 - \frac{n_3}{N}\delta_2\\
\mu_3 - \mu =&(a+\delta_2) - \left(a - \frac{n_1}{N}\delta_1 + \frac{n_3}{N}\delta_2\right)\\
=&\delta_2 + \frac{n_1}{N}\delta_1 - \frac{n_3}{N}\delta_2.
\end{aligned}
$$

Hence, the non-centrality parameter $\lambda$ depends only on $\delta_1$ and $\delta_2$, but not depends on $a$. Therefore, we can set $a = 0$ when we do programming. That is, we can set

$$
\begin{aligned}
&\mu_1 = -\delta_1 \text{ (for mutation homozygotes)},\\
&\mu_2 = 0 \text{ (for heterozygotes)},\\
&\mu_3 = \delta_2 \text{ (for wildtype homozygote)}.
\end{aligned}
\tag{10}
$$

Next, we simplify the expression of $\lambda$.

Denote

$$p = \theta, q = 1 - p.$$

Then we have

$$w_1 = p^2, w_2 = 2pq, w_3 = q^2.$$

The overall mean can be rewritten as

$$\begin{aligned} \mu =& w_1\mu_1 + w_2\mu_2 + w_3\mu 3 \\ =& p^2(-\delta_1) + q^2\delta_2 \\ =& q^2\delta_2 - p^2\delta_1. \end{aligned} \tag{11}$$

Denote

$$\xi = \sum_{i=1}^{3} w_i \left(\mu_i - \mu\right)^2.$$

We can get

$$\begin{aligned} \xi =& \sum_{i=1}^{3} w_i \left(\mu_i^2 - 2\mu_i\mu + \mu^2\right) \\ =& \sum_{i=1}^{3} \left[w_i\mu_i^2 - 2w_i\mu_i\mu + w_i\mu^2\right] \\ =& \sum_{i=1}^{3} w_i\mu_i^2 - 2\mu \sum_{i=1}^{3} w_i\mu_i + \mu^2 \sum_{i=1}^{3} w_i \\ =& \sum_{i=1}^{3} w_i\mu_i^2 - 2\mu^2 + \mu^2 \\ =& \sum_{i=1}^{3} w_i\mu_i^2 - \mu^2 \\ =& w_1\mu_1^2 + w_3\mu_3^2 - \mu^2 \\ =& p^2\delta_1^2 + q^2\delta_2^2 - \left(q^2\delta_2 - p^2\delta_1\right)^2 \\ =& p^2\delta_1^2 + q^2\delta_2^2 - \left(q^4\delta_2^2 + p^4\delta_1^2 - 2p^2q^2\delta_1\delta_2\right) \\ =& \left(p^2 - p^4\right)\delta_1^2 + \left(q^2 - q^4\right)\delta_2^2 + 2p^2q^2\delta_1\delta_2 \\ =& p^2(1-p)(1+p)\delta_1^2 + q^2(1-q)(1+q)\delta_2^2 + 2p^2q^2\delta_1\delta_2 \\ =& p^2q(1+p)\delta_1^2 + q^2p(1+q)\delta_2^2 + 2p^2q^2\delta_1\delta_2. \end{aligned}$$

Hence,

$$\lambda = \frac{N}{\sigma^2}\left[p^2 q(1+p)\delta_1^2 + q^2 p(1+q)\delta_2^2 + 2p^2q^2\delta_1\delta_2\right]. \tag{12}$$

### 2 Power Calculation for Bulk Tissue eQTL Based on SLR

#### 2.1 Introduction

If we would like to test linear relationship between genotype of a SNP and expression of a gene, we can use simple linear regression:

$$\begin{aligned} y_i =& \beta_0 + \beta_1 x_i + \epsilon_i,\ \epsilon_i \sim \mathrm{N}(0, \sigma^2), \\ i =& 1, \ldots, n, \end{aligned} \tag{13}$$

where $y_i$ is the expression level of the gene for the subject $i$ and $x_i$ is the genotype of the $i$-th subject by using additive coding. That is, $x_i = 0$ for wild-type homozygote (containing zero minor allele), $x_i = 1$ for heterozygote (containing one minor allele), and $x_i = 2$ for mutation homozygote (containing two minor allele).

#### 2.2 Mean and variance of additive coded genotype

Denote $A$ as major allele and $a$ as minor allele. Denote $X$ as the random variable recoding the additive coded genotype and denote the genotype frequencies as

$$\begin{aligned} p_0 =& Pr(AA) = Pr(X=0), \\ p_1 =& Pr(Aa) = Pr(X=1), \\ p_2 =& Pr(aa) = Pr(X=2). \end{aligned} \tag{14}$$

Then we can calculate the mean and variance of the genotype

$$\begin{aligned}
\mu_x =& E(X) \\
=& 0 \cdot Pr(X=0) + 1 \cdot Pr(X=1) + 2 \cdot Pr(X=2) \\
=& p_1 + 2p_2, \\
\sigma_x^2 =& E(X^2) - [E(X)]^2 \\
=& \left[0^2 \cdot Pr(X=0) + 1^2 \cdot Pr(X=1) + 2^2 \cdot Pr(X=2)\right] - \mu_x^2 \\
=& p_1 + 4p_2 - \mu_x^2.
\end{aligned} \tag{15}$$

Equivalently,

$$\begin{aligned}
p_1 =& 2\mu_x - \mu_x^2 - \sigma_x^2, \\
p_2 =& \frac{(\sigma_x^2 - \mu_x + \mu_x^2)}{2}.
\end{aligned} \tag{16}$$

##### 2.2.1 Hardy-Weinberg equilibrium

Denote the minor allele frequency (MAF) as $\theta = Pr(a)$. Under Hardy-Weinberg equilibrium, we have the following results:

$$\begin{aligned}
p_0 =& Pr(AA) = (1-\theta)^2, \\
p_1 =& Pr(Aa) = 2\theta(1-\theta), \\
p_2 =& Pr(aa) = \theta^2.
\end{aligned} \tag{17}$$

In this case, we have

$$\begin{aligned}
\mu_x =& 2\theta, \\
\sigma_x^2 =& 2(1-\theta)\theta,
\end{aligned} \tag{18}$$

and

$$\sigma_x^2 = \mu_x - \mu_x^2/2. \tag{19}$$

#### 2.3 Power Calculation of eQTL based on simple linear regression

##### 2.3.1 Power calculation for simple linear regression

The exact power calculation formula derived in this section is an improvement of the appro-lambdamate power calculation formula derived in Dupont and Plummer (1998). For simple linear regression (13), the estimate $\hat{\beta}_1$ of slope is

$$\hat{\beta}_1 = \frac{\sum_{i=1}^{n}(x_i - \bar{x})(y_i - \bar{y})}{\sum_{i=1}^{n}(x_i - \bar{x})^2}, \tag{20}$$

where

$$\bar{x} = \frac{1}{n}\sum_{i=1}^{n} x_i,\ \ \bar{y} = \frac{1}{n}\sum_{i=1}^{n} y_i.$$

Denote

$$L_{xx} = \sum_{i=1}^{n}(x_i - \bar{x})^2 = \sum_{i=1}^{n} x_i^2 - n(\bar{x})^2$$
$$L_{xy} = \sum_{i=1}^{n}(x_i - \bar{x})(y_i - \bar{y}) = \sum_{i=1}^{n} x_i y_i - n\bar{x}\bar{y}.$$

Then

$$\hat{\beta}_1 = \frac{L_{xy}}{L_{xx}}.$$

We are interested in testing the hypotheses:

$$H_0 : \beta_1 = 0 \text{ versus } H_1 : \beta_1 = \delta(\delta \neq 0). \tag{21}$$

Under the alternative hypothesis $H_1$,

$$\hat{\beta}_1 \sim \mathrm{N}\left(\delta,\ \frac{\sigma^2}{L_{xx}}\right). \tag{22}$$

We can construct the following test statistic to test if $\beta_1 = 0$:

$$t = \frac{\hat{\beta}_1}{\sqrt{\hat{\sigma}^2 / L_{xx}}},$$

where

$$\hat{\sigma}^2 = \frac{1}{n-2}\sum_{i=1}^{n}(y_i - \hat{y}_i)^2$$

is the unbiased estimate of $\sigma^2$ (i.e., $E\left(\hat{\sigma}^2\right) = \sigma^2$), and

$$\hat{y}_i = \hat{\beta}_0 + \hat{\beta}_1 x_i$$

and

$$\hat{\beta}_0 = \bar{y} - \hat{\beta}_1 \bar{x}.$$

It can be shown that

$$(n-2)\frac{\hat{\sigma}^2}{\sigma^2} \sim \chi^2_{n-2}. \tag{23}$$

Under $H_0 : \beta_1 = 0$,

$$\frac{\hat{\beta}_1}{\sqrt{\sigma^2/L_{xx}}} \sim N(0,1).$$

It can be shown that $\hat{\beta}_1$ is independent of $\hat{\sigma}^2$.

Note that if $Z \sim N(0,1)$, $V \sim \chi^2_\nu$, and $Z$ and $V$ independent, then $Z/\sqrt{V/\nu} \sim t_\nu$, where $t_\nu$ is the t distribution with $\nu$ degrees of freedom. Hence, we have

$$t = \frac{\hat{\beta}_1}{\sqrt{\hat{\sigma}^2/L_{xx}}} = \frac{\frac{\hat{\beta}_1}{\sqrt{\sigma^2/L_{xx}}}}{\sqrt{(n-2)\frac{\hat{\sigma}^2}{\sigma^2}/(n-2)}} \overset{H_0}{\sim} t_{n-2}.$$

Hence, the Type I error rate $\alpha$ for two-sided test is

$$\alpha = Pr\left(|t| > t_{n-2}(\alpha/2) | H_0\right), \tag{24}$$

where $t_{n-2}(\alpha/2)$ is the upper $100(\alpha/2)\%$ percentile of the $t$ distribution with degree of freedom $n-2$.

We use the following fact: if $Z \sim N(0,1)$, $V \sim \chi^2_\nu$, and $Z$ and $V$ are independent, then

$$\frac{Z+\lambda}{\sqrt{V/\nu}} \sim t_{\nu,\lambda},$$

where $t_{\nu,\lambda}$ is the non-central t distribution with $\nu$ degrees of freedom and non-centrality parameter $\lambda$.

Hence, based on (22), we have

$$\frac{\hat{\beta}_1}{\sqrt{\sigma^2/L_{xx}}} \overset{H_1}{\sim} N\left(\lambda, 1\right),$$

where

$$\lambda = \frac{\delta}{\sqrt{\sigma^2/L_{xx}}} = \frac{\delta}{\sqrt{\sigma^2/\left[(n-1)\tilde{\sigma}_x^2\right]}}, \tag{25}$$

and $\tilde{\sigma}_x^2$ is an unbiased estimate of $\sigma_x^2$:

$$\tilde{\sigma}_x^2 = \frac{1}{n-1} \sum_{i=1}^{n} (x_i - \bar{x})^2 .$$

Under $H_1$ we have

$$t = \frac{\hat{\beta}_1}{\sqrt{\hat{\sigma}^2/L_{xx}}} = \frac{\frac{\hat{\beta}_1}{\sqrt{\sigma^2/L_{xx}}}}{\sqrt{(n-2)\frac{\hat{\sigma}^2}{\sigma^2}/(n-2)}} \overset{H_1}{\sim} t_{n-2,\lambda}. \tag{26}$$

Hence, the exact power is calculated as

$$\begin{aligned} 1-\beta =& Pr\left(|t| > t_{n-2}(\alpha/2)|H_1\right) \\ =& Pr\left(t > t_{n-2}(\alpha/2)|H_1\right) + Pr\left(t < -t_{n-2}(\alpha/2)|H_1\right) \\ =& 1 - T_{n-2,\lambda}\left[t_{n-2}(\alpha/2)\right] + T_{n-2,\lambda}\left[-t_{n-2}(\alpha/2)\right], \end{aligned} \tag{27}$$

where $T_{n-2,\lambda}(a)$ is the value at $a$ of CDF of non-central t distribution with $(n-2)$ degrees of freedom and non-centrality parameter $\lambda$.

Formula (25) shows that $\lambda$ depends on the standard deviation $\sigma$ of random error, which is not easy to estimate or to set its value in design stage of a study. Instead, the standard deviation of the outcome $\sigma_y$ is relative easier to estimate or set based on historical data.

Formulas (1) and (2) in Dupont and Plummer (1998)[1] describe the relationships among the slope $\beta_1$, the variance of the outcome $\sigma_y^2$, the variance of predictor $\sigma_x^2$, and the variance of the random error $\sigma^2$:

$$\sigma^2 = \sigma_y^2 - \beta_1^2 \sigma_x^2. \tag{28}$$

Hence, we can rewrite the non-centrality parameter $\lambda$ as

$$\lambda = \frac{\delta}{\sqrt{\left(\sigma_y^2 - \delta^2 2(1-\hat{\theta})\hat{\theta}\right) / \left[(n-1)2(1-\hat{\theta})\hat{\theta}\right]}}. \tag{29}$$

Since $\sigma^2 > 0$, we require

$$\sigma_y^2 - \delta^2 2(1-\theta)\theta > 0.$$

We can get

$$-\frac{\sigma_y}{\sqrt{2\theta(1-\theta)}} < \delta < \frac{\sigma_y}{\sqrt{2\theta(1-\theta)}}. \tag{30}$$

and

$$\left(\theta - \frac{1}{2}\right)^2 > \frac{1}{4} - \frac{\sigma_y^2}{2\delta^2}.$$

If $\frac{1}{4} - \frac{\sigma_y^2}{2\delta^2} > 0$, we then require

$$\left(\theta - \frac{1}{2}\right) > \sqrt{\frac{1}{4} - \frac{\sigma_y^2}{2\delta^2}}$$

or

$$\left(\theta - \frac{1}{2}\right) < -\sqrt{\frac{1}{4} - \frac{\sigma_y^2}{2\delta^2}}$$

Since $\theta < 0.5$, we require

$$0 < \theta < \frac{1}{2} - \sqrt{\frac{1}{4} - \frac{\sigma_y^2}{2\delta^2}}. \tag{31}$$

### 3 Power Calculation for Single-Cell eQTL

#### 3.1 A Linear Mixed Effects Model

We are interested in testing if a SNP is associated with the expression of a gene based on single cell RNAseq data, which contain $n$ subjects. For each subject, we obtain $m$ cells. For each cell, we measured the expression of $G$ genes.

We assume the following linear mixed effects model to characterize the association between genotype of a given SNP and expression of a given gene:

$$\begin{aligned} y_{ij} =& \beta_{0i} + \beta_1 x_i + \epsilon_{ij}, \\ \beta_{0i} \sim& N\left(\beta_0, \sigma_\beta^2\right), \\ \epsilon_{ij} \sim& N\left(0, \sigma^2\right), \\ i =& 1, \ldots, n, \\ j =& 1, \ldots, m, \end{aligned} \tag{32}$$

where the random intercepts $\beta_{0i}$ and the random error terms $\epsilon_{ij}$ are independent, $n$ is the number of subjects, $m$ is the number of cells per subject, $y_{ij}$ is the gene expression of the $j$-th cell for the $i$-th subject, and $x_i$ is the genotype for the $i$-th subject. $x_i = 0$ indicates that the $i$-th subject is a wildtype homozygote, $x_i = 1$ indicates that the $i$-th subject is a heterozygote, and $x_i = 2$ indicates that the $i$-th subject is a mutation homozygote.

Note that the random intercept $\beta_{0i}$ helps incorporate intra-class correlation between $y_{ij}$ and $y_{ik}$, for $j \neq k$. The covariance between $y_{ij}$ and $y_{ik}$ is

$$\begin{aligned} Cov\left(y_{ij}, y_{ik}\right) =& Cov\left(\beta_{0i} + \beta_1 x_i + \epsilon_{ij}, \beta_{0i} + \beta_1 x_i + \epsilon_{ik}\right) \\ =& Var\left(\beta_{0i}\right) \\ =& \sigma_\beta^2. \end{aligned}$$

#### 3.2 Hypotheses

The mean gene expression for the 3 genotypes are

$$\begin{aligned} E\left(y_{ij}\right) =& \beta_0 \quad \text{if subject } i \text{ is a wildtype homozygote,} \\ E\left(y_{ij}\right) =& \beta_0 + \beta_1 \quad \text{if subject } i \text{ is a heterozygote,} \\ E\left(y_{ij}\right) =& \beta_0 + 2\beta_1 \quad \text{if subject } i \text{ is a mutation homozygote.} \end{aligned}$$

If the slope $\beta_1 = 0$, then all three genotypes have the same mean gene expression $E\left(y_{ij}\right) = \beta_0$. Hence, to test if a SNP is associated with a gene is equivalent to test if the slope $\beta_1 = 0$ or not.

We would like to test the following null hypothesis ($H_0$) and alternative hypothesis ($H_1$):

$$\begin{aligned} H_0 :& \beta_1 = 0, \\ H_1 :& \beta_1 = \delta, \end{aligned} \tag{33}$$

where $\delta \neq 0$.

#### 3.3 Power calculation formula

For a given SNP-gene pair, we derived the power calculation formula for testing Hypotheses (33) as shown below:

$$power = 1 - \Phi\left(z_{\alpha/2} - \frac{\hat{\sigma}_x}{\sigma_y}\frac{\delta\sqrt{m(n-1)}}{\sqrt{1+(m-1)\rho}}\right) + \Phi\left(-z_{\alpha/2} - \frac{\hat{\sigma}_x}{\sigma_y}\frac{\delta\sqrt{m(n-1)}}{\sqrt{1+(m-1)\rho}}\right), \tag{34}$$

where $\alpha$ is the type I error rate, $z_{\alpha/2}$ is the upper $100\alpha/2$ percentile of the standard normal distribution $N(0,1)$, $\sigma_y = \sqrt{\sigma_\beta^2 + \sigma^2}$ is the standard deviation of $y_{ij}$ (i.e., $\sigma_y = \sqrt{Var\left(y_{ij}\right)}$), $\hat{\sigma}_x = \sqrt{\sum_{i=1}^n \left(x_i - \bar{x}\right)^2 / (n-1)}$ is the sample standard deviation of the predictor (i.e., genotype) $x_i$, and $\rho = \sigma_\beta^2 / \left(\sigma_\beta^2 + \sigma^2\right)$ is the intra-class correlation (i.e., correlation between $y_{ij}$ and $y_{ik}$).

The power calculation formula for testing Hypotheses (33) for genotypes under Hardy-Weinberg Equilibrium is:

$$\begin{aligned} power =& 1 - \Phi\left(z_{\alpha/2} - \frac{\sqrt{2\theta\left(1-\theta\right)}}{\sigma_y} \frac{\delta\sqrt{m(n-1)}}{\sqrt{1+(m-1)\rho}}\right) \\ &+ \Phi\left(-z_{\alpha/2} - \frac{\sqrt{2\theta\left(1-\theta\right)}}{\sigma_y} \frac{\delta\sqrt{m(n-1)}}{\sqrt{1+(m-1)\rho}}\right). \end{aligned} \tag{35}$$

where $\theta$ is the minor allele frequency of the SNP.

The details of the derivations are shown in Appendices.

#### 3.4 Discussion

Note that we assume the gene expression levels $y_{ij}$ are normally distributed. However, RNAseq data are counts and many counts are zero. So $y_{ij}$ could not be normally distributed. We might use data transformations (e.g., R package *countTransformers*[4] developed in Zhang et al., 2019[5]). However, usually we can only make sure the median and mean are close, while the transformed data are still not normally distributed. The effect of using normal assumption to fit non-normal data is false-positive inflation. So the power calculated will be higher than the true power. In future, we will derive power calculation formula based on mixed effects negative binomial regressions that are popular in fitting RNAseq data. The challenge is that no closed-form power calculation formulas can be derived for negative binomial regressions, let alone mixed effects negative binomial regressions. We will try to derive approximate power-calculation formulas and to use simulation approach.

### Appendix

#### A Vector representation

In this section, we derive the vector representations of Model (32).

We represent $\beta_{0i}$ by

$$\beta_{0i} = \beta_0 + \xi_i,$$

where

$$\xi_i \sim N\left(0, \sigma_\beta^2\right).$$

Denote

$$e_{ij} = \xi_i + \epsilon_{ij}.$$

Then

$$e_{ij} \sim N\left(0, \sigma_\beta^2 + \sigma^2\right).$$

Denote

$$\sigma_y^2 = \sigma_\beta^2 + \sigma^2.$$

Note that

$$Var\left(y_{ij}\right) = Var\left(\beta_{0i}\right) + Var\left(\epsilon_{ij}\right) = \sigma_\beta^2 + \sigma^2 = \sigma_y^2. \tag{A1}$$

Model (32) can be rewritten as

$$\begin{aligned} y_{ij} =& \beta_0 + \beta_1 x_i + e_{ij}, \\ e_{ij} \sim& N\left(0, \sigma_y^2\right), \\ i =& 1, \dots, n, \\ j =& 1, \dots, m, \end{aligned} \tag{A2}$$

with $Cov\left(y_{ij}, y_{ik}\right) = \sigma_\beta^2$ for $j \neq k$.

Denote

$$\boldsymbol{y}_i = \begin{pmatrix} y_{i1} \\ \vdots \\ y_{im} \end{pmatrix}, \quad \boldsymbol{e}_i = \begin{pmatrix} e_{i1} \\ \vdots \\ e_{im} \end{pmatrix}, \quad \boldsymbol{u}_i = \begin{pmatrix} 1 \\ x_i \end{pmatrix}, \quad \boldsymbol{\beta} = \begin{pmatrix} \beta_0 \\ \beta_1 \end{pmatrix}.$$

Also, denote $\mathbf{1}_m$ as the $m \times 1$ vector of ones, $\mathbf{0}_m$ as the $m \times 1$ vector of zeros, and $\boldsymbol{I}_m$ as the

$m \times m$ identity matrix.

Model (A2) can be rewritten as

$$\begin{aligned} \boldsymbol{y}_i =& \mathbf{1}_m \boldsymbol{u}_i^T \boldsymbol{\beta} + \boldsymbol{e}_i, \\ \boldsymbol{e}_i \sim& N\left(\mathbf{0}_m, \boldsymbol{\Sigma}\right), \\ i =& 1, \ldots, n, \end{aligned} \tag{A3}$$

where $\boldsymbol{y}_i$ is a $m \times 1$ vector, $\mathbf{1}_m$ is a $m \times 1$ vector, $\boldsymbol{u}_i^T$ is a $1 \times 2$ vector, $\boldsymbol{\beta}$ is a $2 \times 1$ vector, $\boldsymbol{e}_i$ is a $m \times 1$ vector, $\boldsymbol{\Sigma}$ is a $m \times m$ matrix

$$\boldsymbol{\Sigma} = \begin{pmatrix} \sigma_y^2 & & \sigma_\beta^2 \\ & \ddots & \\ \sigma_\beta^2 & & \sigma_y^2 \end{pmatrix} = \sigma_y^2 \begin{pmatrix} 1 & & \rho \\ & \ddots & \\ \rho & & 1 \end{pmatrix},$$

and

$$\rho = \frac{\sigma_\beta^2}{\sigma_y^2} = \frac{\sigma_\beta^2}{\sigma_\beta^2 + \sigma^2}.$$

Note that $\rho$ is also called intra-class correlation (ICC).

Denote

$$\boldsymbol{R} = \begin{pmatrix} 1 & & \rho \\ & \ddots & \\ \rho & & 1 \end{pmatrix}.$$

Then

$$\boldsymbol{\Sigma} = \sigma_y^2 \boldsymbol{R}.$$

Denote

$$\boldsymbol{y} = \begin{pmatrix} \boldsymbol{y}_1 \\ \vdots \\ \boldsymbol{y}_n \end{pmatrix}, \quad \boldsymbol{U} = \begin{pmatrix} \mathbf{1}_m \boldsymbol{u}_1^T \\ \vdots \\ \mathbf{1}_m \boldsymbol{u}_n^T \end{pmatrix}, \quad \boldsymbol{e} = \begin{pmatrix} \boldsymbol{e}_1 \\ \vdots \\ \boldsymbol{e}_n \end{pmatrix}, \quad \boldsymbol{\Omega} = \begin{pmatrix} \boldsymbol{\Sigma} & & \mathbf{0} \\ & \ddots & \\ \mathbf{0} & & \boldsymbol{\Sigma} \end{pmatrix}.$$

Model (A3) can be rewritten as

$$\begin{aligned} \boldsymbol{y} =& \boldsymbol{U} \boldsymbol{\beta} + \boldsymbol{e}, \\ \boldsymbol{e} \sim& N\left(\mathbf{0}_{(n \times m) \times 1}, \boldsymbol{\Omega}\right), \end{aligned} \tag{A4}$$

where $\boldsymbol{y}$ is a $(n \times m) \times 1$ vector, $\boldsymbol{U}$ is a $(n \times m) \times 2$ matrix, $\boldsymbol{\beta}$ is a $2 \times 1$ vector, and $\boldsymbol{e}$ is a $(n \times m) \times 1$ vector.

### B Generalized least squares estimate when variance-covariance matrix is known

Denote the weighted distance between $\boldsymbol{y}$ and the linear part $\boldsymbol{U\beta}$ in Model (A4) as

$$\begin{aligned} g(\boldsymbol{\beta}) &= (\boldsymbol{y} - \boldsymbol{U\beta})^T \boldsymbol{\Omega}^{-1} (\boldsymbol{y} - \boldsymbol{U\beta}) \\ &= \boldsymbol{y}^T \boldsymbol{\Omega}^{-1} \boldsymbol{y} - 2\boldsymbol{y}^T \boldsymbol{\Omega}^{-1} \boldsymbol{U\beta} + \boldsymbol{\beta}^T \boldsymbol{U}^T \boldsymbol{\Omega}^{-1} \boldsymbol{U\beta}. \end{aligned} \tag{A5}$$

The generalized least squares estimate of $\boldsymbol{\beta}$ when the variance-covariance matrix $\boldsymbol{\Omega}$ is known is obtained by solving the following minimization problem:

$$min_{\boldsymbol{\beta}} g(\boldsymbol{\beta}). \tag{A6}$$

The first order partial derivative of $g(\boldsymbol{\beta})$ to $\boldsymbol{\beta}$ is

$$\frac{\partial g(\boldsymbol{\beta})}{\partial \boldsymbol{\beta}} = -2\boldsymbol{U}^T \boldsymbol{\Omega}^{-1} \boldsymbol{y} + 2\boldsymbol{U}^T \boldsymbol{\Omega}^{-1} \boldsymbol{U\beta}.$$

Let the first partial derivative be equal to zero, we can get

$$\hat{\boldsymbol{\beta}} = \left(\boldsymbol{U}^T \boldsymbol{\Omega}^{-1} \boldsymbol{U}\right)^{-1} \boldsymbol{U}^T \boldsymbol{\Omega}^{-1} \boldsymbol{y}. \tag{A7}$$

The second order partial derivative of $g(\boldsymbol{\beta})$ to $\boldsymbol{\beta}$ is

$$\frac{\partial^2 g(\boldsymbol{\beta})}{\partial \boldsymbol{\beta} \partial \boldsymbol{\beta}^T} = 2\boldsymbol{U}^T \boldsymbol{\Omega}^{-1} \boldsymbol{U},$$

which is positive definite. Hence, $\hat{\boldsymbol{\beta}}$ minimizes, not maximize, the objective function $g(\boldsymbol{\beta})$. We call $\hat{\boldsymbol{\beta}}$ the generalized least squares estimate of $\boldsymbol{\beta}$ when $\boldsymbol{\Omega}$ is known.

#### C Mean, variance, and distribution of the generalized least squares estimate

The mean of $\hat{\boldsymbol{\beta}}$ is

$$\begin{aligned} E\left(\hat{\boldsymbol{\beta}}\right) &= \left(\boldsymbol{U}^T\boldsymbol{\Omega}^{-1}\boldsymbol{U}\right)^{-1}\boldsymbol{U}^T\boldsymbol{\Omega}^{-1}E\left(\boldsymbol{y}\right) \\ &= \left(\boldsymbol{U}^T\boldsymbol{\Omega}^{-1}\boldsymbol{U}\right)^{-1}\boldsymbol{U}^T\boldsymbol{\Omega}^{-1}\boldsymbol{U}\boldsymbol{\beta} \\ &= \boldsymbol{\beta}. \end{aligned}$$

That is $\hat{\boldsymbol{\beta}}$ is an unbiased estimate of $\boldsymbol{\beta}$.

The variance-covariance matrix of $\hat{\boldsymbol{\beta}}$ is

$$\begin{aligned} Cov\left(\hat{\boldsymbol{\beta}}\right) &= \left(\boldsymbol{U}^T\boldsymbol{\Omega}^{-1}\boldsymbol{U}\right)^{-1}\boldsymbol{U}^T\boldsymbol{\Omega}^{-1}Cov\left(\boldsymbol{y}\right)\boldsymbol{\Omega}^{-1}\boldsymbol{U}\left(\boldsymbol{U}^T\boldsymbol{\Omega}^{-1}\boldsymbol{U}\right)^{-1} \\ &= \left(\boldsymbol{U}^T\boldsymbol{\Omega}^{-1}\boldsymbol{U}\right)^{-1}\boldsymbol{U}^T\boldsymbol{\Omega}^{-1}\boldsymbol{\Omega}\boldsymbol{\Omega}^{-1}\boldsymbol{U}\left(\boldsymbol{U}^T\boldsymbol{\Omega}^{-1}\boldsymbol{U}\right)^{-1} \\ &= \left(\boldsymbol{U}^T\boldsymbol{\Omega}^{-1}\boldsymbol{U}\right)^{-1}. \end{aligned} \tag{A8}$$

Since $\boldsymbol{y}$ is normally distributed, hence $\hat{\boldsymbol{\beta}}$ is also normally distributed.

#### D Calculating the power for testing if the slope is equal to zero

To test the null hypothesis $H_0 : \beta_1 = 0$ versus the alternative hypothesis $H_1 : \beta_1 = \delta$, where $\delta \neq 0$, we can construct the test statistic

$$Z = \frac{\hat{\beta}_1}{\sqrt{Var\left(\hat{\beta}_1\right)}}. \tag{A9}$$

Note that we assume $\boldsymbol{\Omega}$ is known. Hence, $Z$ is a statistic. Otherwise (i.e., if $\boldsymbol{\Omega}$ is unknown), $Z$ is not a statistic since it contains unknown parameters.

Under $H_0 : \beta_1 = 0$, $Z$ follows standard normal distribution $N(0, 1)$. Under $H_1 : \beta_1 = \delta$, where $\delta \neq 0$,

$$\frac{\hat{\beta}_1 - \delta}{\sqrt{Var\left(\hat{\beta}_1\right)}} \sim N(0,1).$$

If we set the decision rule as

$$\text{reject } H_0 : \beta_1 = 0 \text{ if } |Z| > z_{\alpha/2},$$

then the Type I error rate is equal to $\alpha$, where $z_{\alpha/2}$ is the upper $100\alpha/2$ percentile. That is,

$$Pr\left(|Z| > z_{\alpha/2} | H_0\right) = \alpha$$

We also can calculate the power for testing he null hypothesis $H_0 : \beta_1 = 0$ versus the alternative hypothesis $H_1 : \beta_1 = \delta$, where $\delta \neq 0$.

$$\begin{aligned} power =& Pr\left(|Z| > z_{\alpha/2} | H_1\right) \\ =& Pr\left(Z > z_{\alpha/2} | H_1\right) + Pr\left(Z < -z_{\alpha/2} | H_1\right). \end{aligned}$$

We can get

$$\begin{aligned} & Pr\left(Z > z_{\alpha/2} | H_1\right) \\ =& Pr\left(\left.\frac{\hat{\beta}_1}{\sqrt{Var\left(\hat{\beta}_1\right)}} > z_{\alpha/2} \right| H_1\right) \\ =& Pr\left(\left.\frac{\hat{\beta}_1 - \delta + \delta}{\sqrt{Var\left(\hat{\beta}_1\right)}} > z_{\alpha/2} \right| H_1\right) \\ =& Pr\left(\left.\frac{\hat{\beta}_1 - \delta}{\sqrt{Var\left(\hat{\beta}_1\right)}} > z_{\alpha/2} - \frac{\delta}{\sqrt{Var\left(\hat{\beta}_1\right)}} \right| H_1\right) \\ =& 1 - \Phi\left(z_{\alpha/2} - \frac{\delta}{\sqrt{Var\left(\hat{\beta}_1\right)}}\right), \end{aligned}$$

where $\Phi$ is the cumulative distribution function of the standard normal distribution $N(0,1)$.

Similarly, we can get

$$
\begin{aligned}
& Pr\left(Z < -z_{\alpha/2} | H_1\right) \\
=& Pr\left(\left.\frac{\hat{\beta}_1}{\sqrt{Var\left(\hat{\beta}_1\right)}} < -z_{\alpha/2}\right| H_1\right) \\
=& Pr\left(\left.\frac{\hat{\beta}_1 - \delta + \delta}{\sqrt{Var\left(\hat{\beta}_1\right)}} < -z_{\alpha/2}\right| H_1\right) \\
=& Pr\left(\left.\frac{\hat{\beta}_1 - \delta}{\sqrt{Var\left(\hat{\beta}_1\right)}} < -z_{\alpha/2} - \frac{\delta}{\sqrt{Var\left(\hat{\beta}_1\right)}}\right| H_1\right) \\
=& \Phi\left(-z_{\alpha/2} - \frac{\delta}{\sqrt{Var\left(\hat{\beta}_1\right)}}\right).
\end{aligned}
$$

Therefore the power is

$$
power = 1 - \Phi\left(z_{\alpha/2} - \frac{\delta}{\sqrt{Var\left(\hat{\beta}_1\right)}}\right) + \Phi\left(-z_{\alpha/2} - \frac{\delta}{\sqrt{Var\left(\hat{\beta}_1\right)}}\right). \tag{A10}
$$

### E Calculation of the variance of the slope estimate

Based on Formula (A8), we have

$$
Cov\left(\hat{\boldsymbol{\beta}}\right) = \left(\boldsymbol{U}^T \boldsymbol{\Omega}^{-1} \boldsymbol{U}\right)^{-1}.
$$

We first calculate $\mathbf{\Omega}^{-1}$. We have

$$\mathbf{\Omega}^{-1} = \begin{pmatrix} \mathbf{\Sigma}^{-1} & & \mathbf{0} \\ & \ddots & \\ \mathbf{0} & & \mathbf{\Sigma}^{-1} \end{pmatrix}.$$

We then calculate $\mathbf{\Sigma}^{-1}$. We have

$$\mathbf{\Sigma}^{-1} = \frac{1}{\sigma_y^2}\boldsymbol{R}^{-1}.$$

We next calculate $\boldsymbol{R}^{-1}$. We can rewrite $\boldsymbol{R}$ to

$$\boldsymbol{R} = (1-\rho)\left(\boldsymbol{I}_m + \frac{\rho}{1-\rho}\mathbf{1}_m\mathbf{1}_m^T\right).$$

Then we have

$$\boldsymbol{R}^{-1} = \frac{1}{(1-\rho)}\left(\boldsymbol{I}_m + \frac{\rho}{1-\rho}\mathbf{1}_m\mathbf{1}_m^T\right)^{-1}.$$

Based on the matrix theories, we have the following results

$$\left(\boldsymbol{A} + \boldsymbol{b}\boldsymbol{c}^T\right)^{-1} = \boldsymbol{A}^{-1} - \frac{\boldsymbol{A}^{-1}\boldsymbol{b}\boldsymbol{c}\boldsymbol{A}^{-1}}{1 + \boldsymbol{c}^T\boldsymbol{A}^{-1}\boldsymbol{b}},$$

where $\boldsymbol{A}$ is a matrix, $\boldsymbol{b}$ and $\boldsymbol{c}$ are vectors.

Let $\boldsymbol{A} = \boldsymbol{I}_m$, $\boldsymbol{b} = \frac{\rho}{1-\rho}\mathbf{1}_m$, $\boldsymbol{c} = \mathbf{1}_m$. We have $\boldsymbol{A}^{-1} = \boldsymbol{I}_m$, $\mathbf{1}_m^T\mathbf{1}_m = m$, and

$$\begin{aligned}&\left(\boldsymbol{I}_m + \frac{\rho}{1-\rho}\mathbf{1}_m\mathbf{1}_m^T\right)^{-1} \\ =&\boldsymbol{I}_m - \frac{\frac{\rho}{1-\rho}\mathbf{1}_m\mathbf{1}_m^T}{1 + \frac{\rho}{1-\rho}m} \\ =&\boldsymbol{I}_m - \frac{\rho\mathbf{1}_m\mathbf{1}_m^T}{(1-\rho) + \rho m} \\ =&\boldsymbol{I}_m - \frac{\rho\mathbf{1}_m\mathbf{1}_m^T}{1 + (m-1)\rho}.\end{aligned}$$

Hence, we have

$$\boldsymbol{R}^{-1} = \frac{1}{(1-\rho)}\left[\boldsymbol{I}_m - \frac{\rho\mathbf{1}_m\mathbf{1}_m^T}{1 + (m-1)\rho}\right]$$

and

$$\boldsymbol{\Sigma}^{-1} = \frac{1}{\sigma_y^2(1-\rho)}\left[\boldsymbol{I}_m - \frac{\rho \mathbf{1}_m \mathbf{1}_m^T}{1+(m-1)\rho}\right]. \tag{A11}$$

Next, we calculate

$$\begin{aligned}
\boldsymbol{U}^T\boldsymbol{\Omega}^{-1}\boldsymbol{U} &= \left(\boldsymbol{u}_1\mathbf{1}_m^T, \ldots, \boldsymbol{u}_n\mathbf{1}_m^T\right)\begin{pmatrix} \boldsymbol{\Sigma}^{-1} & & \mathbf{0} \\ & \ddots & \\ \mathbf{0} & & \boldsymbol{\Sigma}^{-1} \end{pmatrix}\begin{pmatrix} \mathbf{1}_m\boldsymbol{u}_1^T \vdots \\ \mathbf{1}_m\boldsymbol{u}_n^T \end{pmatrix} \\
&= \sum_{i=1}^n \boldsymbol{u}_i\mathbf{1}_m^T\boldsymbol{\Sigma}^{-1}\mathbf{1}_m\boldsymbol{u}_i^T \\
&= \mathbf{1}_m^T\boldsymbol{\Sigma}^{-1}\mathbf{1}_m\sum_{i=1}^n \boldsymbol{u}_i\boldsymbol{u}_i^T.
\end{aligned}$$

We have

$$\begin{aligned}
\mathbf{1}_m^T\boldsymbol{\Sigma}^{-1}\mathbf{1}_m &= \frac{1}{\sigma_y^2(1-\rho)}\left[\mathbf{1}_m^T\mathbf{1}_m - \frac{\rho\mathbf{1}_m^T\mathbf{1}_m\mathbf{1}_m^T\mathbf{1}_m}{1+(m-1)\rho}\right] \\
&= \frac{1}{\sigma_y^2(1-\rho)}\left[m - \frac{\rho m^2}{1+(m-1)\rho}\right] \\
&= \frac{m}{\sigma_y^2(1-\rho)}\left[1 - \frac{\rho m}{1+(m-1)\rho}\right] \\
&= \frac{m}{\sigma_y^2(1-\rho)}\frac{[1+(m-1)\rho-\rho m]}{[1+(m-1)\rho]} \\
&= \frac{m}{\sigma_y^2(1-\rho)}\frac{(1-\rho)}{[1+(m-1)\rho]} \\
&= \frac{m}{\sigma_y^2[1+(m-1)\rho]}.
\end{aligned}$$

We also can get

$$\begin{aligned}
\boldsymbol{u}_i\boldsymbol{u}_i^T &= \begin{pmatrix} 1 \\ x_i \end{pmatrix}(1, x_i) \\
&= \begin{pmatrix} 1 & x_i \\ x_i & x_i^2 \end{pmatrix}.
\end{aligned}$$

Hence, we have

$$
\begin{aligned}
\boldsymbol{U}^T\boldsymbol{\Omega}^{-1}\boldsymbol{U} =& \mathbf{1}_m^T\boldsymbol{\Sigma}^{-1}\mathbf{1}_m\sum_{i=1}^{n}\boldsymbol{u}_i\boldsymbol{u}_i^T \\
=& \frac{m}{\sigma_y^2\left[1+(m-1)\rho\right]}\begin{pmatrix} n & \sum_{i=1}^n x_i \\ \sum_{i=1}^n x_i & \sum_{i=1}^n x_i^2 \end{pmatrix} \\
=& \frac{nm}{\sigma_y^2\left[1+(m-1)\rho\right]}\begin{pmatrix} 1 & \bar{x} \\ \bar{x} & \sum_{i=1}^n x_i^2/n \end{pmatrix}.
\end{aligned}
$$

Note that based on the matrix theories, the inverse of a $2\times 2$ matrix

$$
\boldsymbol{A} = \begin{pmatrix} a & b \\ c & d \end{pmatrix}
$$

is

$$
\boldsymbol{A}^{-1} = \frac{\begin{pmatrix} d & -b \\ -c & a \end{pmatrix}}{ad-bc}.
$$

Therefore,

$$
\begin{aligned}
Cov\left(\hat{\boldsymbol{\beta}}\right) &= \left(\boldsymbol{U}^T\boldsymbol{\Omega}^{-1}\boldsymbol{U}\right)^{-1} \\
&= \frac{\sigma_y^2\left[1+(m-1)\rho\right]}{nm}\frac{1}{\left(\sum_{i=1}^n x_i^2/n-\bar{x}^2\right)}\begin{pmatrix} \sum_{i=1}^n x_i^2/n & -\bar{x} \\ -\bar{x} & 1 \end{pmatrix} \\
&= \frac{\sigma_y^2\left[1+(m-1)\rho\right]}{nm}\frac{1}{\frac{1}{n}\sum_{i=1}^n\left(x_i-\bar{x}\right)^2}\begin{pmatrix} \sum_{i=1}^n x_i^2/n & -\bar{x} \\ -\bar{x} & 1 \end{pmatrix} \\
&= \frac{\sigma_y^2\left[1+(m-1)\rho\right]}{m}\frac{1}{\sum_{i=1}^n\left(x_i-\bar{x}\right)^2}\begin{pmatrix} \sum_{i=1}^n x_i^2/n & -\bar{x} \\ -\bar{x} & 1 \end{pmatrix} \\
&= \frac{\sigma_y^2\left[1+(m-1)\rho\right]}{m(n-1)}\frac{1}{\hat{\sigma}_x^2}\begin{pmatrix} \sum_{i=1}^n x_i^2/n & -\bar{x} \\ -\bar{x} & 1 \end{pmatrix} \\
&= \frac{\left[1+(m-1)\rho\right]}{m(n-1)}\frac{\sigma_y^2}{\hat{\sigma}_x^2}\begin{pmatrix} \sum_{i=1}^n x_i^2/n & -\bar{x} \\ -\bar{x} & 1 \end{pmatrix},
\end{aligned}
\tag{A12}
$$

where

$$
\hat{\sigma}_x^2 = \frac{1}{(n-1)}\sum_{i=1}^{n}\left(x_i-\bar{x}\right)^2.
$$

Finally, we obtain

$$Var\left(\hat{\beta}_1\right) = \frac{[1+(m-1)\rho]}{m(n-1)}\frac{\sigma_y^2}{\hat{\sigma}_x^2}. \tag{A13}$$

Note that $Var\left(\hat{\beta}_1\right)$ is the variance conditional on $x_1$, …, $x_n$.

### F Power calculation formula revisit

Based on Formulas (A10) and (A13), the power calculation formula for testing Hypotheses (33) can be rewritten as:

$$\begin{aligned} power =& 1-\Phi\left(z_{\alpha/2}-\frac{\delta}{\sqrt{Var\left(\hat{\beta}_1\right)}}\right)+\Phi\left(-z_{\alpha/2}-\frac{\delta}{\sqrt{Var\left(\hat{\beta}_1\right)}}\right) \\ =& 1-\Phi\left(z_{\alpha/2}-\frac{\hat{\sigma}_x}{\sigma_y}\frac{\delta\sqrt{m(n-1)}}{\sqrt{1+(m-1)\rho}}\right)+\Phi\left(-z_{\alpha/2}-\frac{\hat{\sigma}_x}{\sigma_y}\frac{\delta\sqrt{m(n-1)}}{\sqrt{1+(m-1)\rho}}\right), \end{aligned}$$

where $\alpha$ is the type I error rate, $z_{\alpha/2}$ is the upper $100\alpha/2$ percentile of the standard normal distribution $N(0,1)$, $\sigma_y = \sqrt{\sigma_\beta^2+\sigma^2}$, $\hat{\sigma}_x = \sqrt{\sum_{i=1}^n\left(x_i-\bar{x}\right)^2/(n-1)}$, and $\rho = \sigma_\beta^2/\left(\sigma_\beta^2+\sigma^2\right)$.

### G Variance of genotype under Hardy-Weinberg Equilibrium

For the given SNP, suppose its minor allele frequency (MAF) is $\theta$ ($0<\theta<0.5$). Then under Hardy-Weinberg Equilibrium, the genotype frequencies are

$$\begin{aligned} Pr\left(x_i=2\right) =& \theta^2, \\ Pr\left(x_i=1\right) =& 2\theta\left(1-\theta\right), \\ Pr\left(x_i=0\right) =& \left(1-\theta\right)^2, \end{aligned}$$

The mean genotype is

$$
\begin{aligned}
E\left(x_i\right) =& 2 \times Pr\left(x_i = 2\right) + 1 \times Pr\left(x_i = 1\right) + 0 \times Pr\left(x_i = 0\right) \\
=& 2\theta^2 + 2\theta\left(1-\theta\right) \\
=& 2\theta.
\end{aligned}
$$

We also can derive the second moment of genotype

$$
\begin{aligned}
E\left(x_i^2\right) =& 2^2 \times Pr\left(x_i = 2\right) + 1^2 \times Pr\left(x_i = 1\right) + 0^2 \times Pr\left(x_i = 0\right) \\
=& 4\theta^2 + 2\theta\left(1-\theta\right) \\
=& 2\theta^2 + 2\theta \\
=& 2\theta\left(1+\theta\right).
\end{aligned}
$$

The variance of genotype is

$$
\begin{aligned}
Var\left(x_i^2\right) =& E\left(x_i^2\right) - \left[E\left(x_i\right)\right]^2 \\
=& 2\theta\left(1+\theta\right) - 4\theta^2 \\
=& 2\theta\left(1-\theta\right).
\end{aligned}
$$

### H Power calculation formula for genotypes under Hardy-Weinberg Equilibrium

Hence, the power calculation formula for testing Hypotheses (33) for genotypes under Hardy-Weinberg Equilibrium is:

$$
\begin{aligned}
power =& 1 - \Phi\left(z_{\alpha/2} - \frac{\sqrt{2\theta\left(1-\theta\right)}}{\sigma_y} \frac{\delta\sqrt{m(n-1)}}{\sqrt{1+(m-1)\rho}}\right) \\
&+ \Phi\left(-z_{\alpha/2} - \frac{\sqrt{2\theta\left(1-\theta\right)}}{\sigma_y} \frac{\delta\sqrt{m(n-1)}}{\sqrt{1+(m-1)\rho}}\right).
\end{aligned}
$$

where $\alpha$ is the type I error rate, $z_{\alpha/2}$ is the upper $100\alpha/2$ percentile of the standard normal distribution $N(0,1)$, $\sigma_y = \sqrt{\sigma_\beta^2 + \sigma^2}$, $\theta$ is the minor allele frequency, and $\rho = \sigma_\beta^2 / \left(\sigma_\beta^2 + \sigma^2\right)$ is the intra-class correlation.
